# Plasmid-based CRISPR-Cas9 gene editing in multiple *Candida* species

**DOI:** 10.1101/557926

**Authors:** Lisa Lombardi, João Oliveira-Pacheco, Geraldine Butler

**Author notes:** These authors contributed equally to the work (Lisa Lombardi, João Oliveira-Pacheco). Corresponding author: Geraldine Butler, School of Biomolecular and Biomedical Science, Conway Institute, University College Dublin, Belfield, Dublin 4, Ireland.

## Abstract

Many *Candida* species that cause infection have diploid genomes, and do not undergo classical meiosis. The application of CRISPR-Cas9 gene editing systems have therefore greatly facilitated the generation of gene disruptions, and the introduction of specific polymorphisms. However, CRISPR methods are not yet available for all *Candida* species. We describe here an adaption of a previously developed CRISPR system in *Candida parapsilosis* that uses an autonomously replicating plasmid. Guide RNAs can be introduced in a single cloning step, and are released by cleavage between a tRNA and a ribozyme. The plasmid also contains *CAS9* and a selectable nourseothricin *SAT1* marker. It can be used for markerless editing in *C. parapsilosis, C. orthopsilosis* and *C. metapsilosis*. We also show that CRISPR can easily be used to introduce molecular barcodes, and to reintroduce wild type sequences into edited strains. Heterozygous mutations can be generated, either by careful selection of the distance between the polymorphism and the Cas9 cut site, or by providing two different repair templates at the same time. In addition, we have constructed a different autonomously replicating plasmid for CRISPR-Cas9 editing in *Candida tropicalis*. We show that editing can easily be carried out in multiple *C. tropicalis* isolates. NHEJ repair occurs at a high level in *C. metapsilosis* and *C. tropicalis*.

**IMPORTANCE:** *Candida* species are a major cause of infection worldwide. The species associated with infection vary with geographical location, and patient population. Infection with *Candida tropicalis* is particularly common in South America and Asia, and *Candida parapsilosis* infections are more common in the very young. Molecular methods for manipulating the genomes of these species are still lacking. We describe a simple and efficient CRISPR-based gene editing system that can be applied in the *C. parapsilosis* species group, including the sister species *Candida orthopsilosis* and *Candida metapsilosis*. We have also constructed a separate system for gene editing in *C. tropicalis*.

## INTRODUCTION

Opportunistic yeast pathogens including *Candida* species cause a wide variety of infections, ranging from superficial to systemic, which can often be fatal (1). Infection of premature neonates, the elderly, and immunocompromised patients is particularly common (2). Extensive use of catheters, broad-spectrum antibiotics, and abdominal surgery also favor the spread of pathogenic yeasts from their normal commensal niches (2).

More than 30 *Candida* species are known to cause disease (3, 4). Although *C. albicans* is the most common cause of candidiasis, the emergence of *non-albicans Candida* species such as *Candida dubliniensis, Candida glabrata, Candida krusei, Candida parapsilosis*, and *Candida tropicalis* has increased over the past decades (5, 6), and *Candida auris* has recently been reported as an emerging multidrug-resistant species (7, 8). Many of the well characterized *Candida* species, including *C. albicans, C. parapsilosis* and *C. tropicalis* belong to the CUG-Ser1 clade, in which the CUG codon is translated as serine rather than leucine (9, 10). *C. tropicalis* is particularly prevalent in in South America and Asia (4). The *C. parapsilosis* group, which includes *Candida orthopsilosis* and *Candida metapsilosis*, are most common in the very young and the very old (11).

*C. albicans, C. tropicalis* and the *C. parapsilosis* species complex have diploid genomes and do not undergo meiosis, which makes generating deletion strains a difficult process. Each allele needs to be targeted independently. Several gene deletion methods based on homologous recombination were developed for *C. albicans*, including sequential replacement of alleles with a recyclable nourseothricin resistance marker (12), or with different markers in an auxotrophic background (13). Some systems were adapted for use in *C. parapsilosis* (14–16) and *C. tropicalis* (17–19). Recently, counter selection against the mazF gene of *E. coli* was used to make markerless disruptions in *C. tropicalis* (20). The recent discovery of haploid forms of *C. albicans* has enabled disruption strategies in this species that are not yet applicable in the others (21–23).

The advent of clustered regularly interspaced short palindromic repeat (CRISPR)-based gene editing tools has revolutionized studies in many *Candida* species, including *C. albicans, C. glabrata, C. lusitaniae* and *C. auris* (7, 24–35). Various approaches have been used, including integrating *CAS9* in the genome (e.g. (24)), transient expression of *CAS9* (31), or providing Cas9 as part of an RNA-protein complex (34, 35). Some systems require cloning (e.g. (25)) and some can be constructed using only PCR (e.g. (30, 31)). Some introduce markers at the target site that can be subsequently removed (e.g. (30)). We recently described a CRISPR-Cas9 system on a replicating plasmid that can be used for markerless gene editing in *C. parapsilosis* (27). The *CAS9* containing plasmid is quickly lost in the absence of selection. The system has also been applied in *C. orthopsilosis* (36). Although the plasmid-based gene editing method is efficient, two cloning steps were required, which means it was not readily applicable to large scale efforts. Here, we adapt the plasmid system so that the guide RNA can be introduced in a single cloning step. The system can also be used for gene editing in *C. orthopsilosis* and *C. metapsilosis*. Furthermore, we made a new plasmid for CRISPR editing in *C. tropicalis*. We show that with careful guide design, CRISPR can be used to generate heterozygous variants.

## Results and discussion

### Modification of plasmid-based CRISPR-Cas9 gene editing in *C. parapsilosis*

We first described a plasmid-based CRISPR-Cas9 editing system in *C. parapsilosis* in 2017 (27). The guide RNA is introduced between two ribozymes (Hammerhead and Hepatitis Delta Virus (HDV)) in a two step cloning process, and is released by self-cleavage of the ribozymes. Changing the guide RNA to target a new gene also requires changing bases in the Hammerhead ribozyme. Here, we replace Hammerhead with a tRNA^Ala^ sequence from *C. parapsilosis* using a synthetic construct, based on systems described by Ng and Dean (37) (pCP-tRNA, Fig. 1A). The gRNA is now introduced in a single step by designing two 20 base oligonucleotides with overhanging ends compatible with two SapI sites (Fig. 1B). The mature sgRNA molecule is released by cleaving after the tRNA^Ala^ by endogenous yeast RNase Z endonuclease, and self-splicing before the HDV ribozyme (Fig. 1D). The plasmid expresses *CAS9* and contains a selectable marker (nourseothricin resistance, Fig. 1A). Plasmid loss is induced after only two passages in the absence of selection (Fig. 1C). Gene editing is carried out in a single transformation step, by introducing the plasmid together with a repair template. Repair templates (described in (27)) are generated by overlapping PCR, and include homology arms of 50 and 34 bp upstream and downstream of the cut site, respectively. Fig. 1E shows editing of *CpADE2* using pCP-tRNA by the introduction of two stop codons, using sgADE2-B as previously described in Lombardi et al. (27). Disruptions in *ADE2* are easily detected because they accumulate a red/pink pigment on YPD media, due to a defect in adenine biosynthesis. Transformants were screened by PCR using one primer specific to the edited site, followed by sequencing (Table S2, doi:10.6084/m9.figshare.7752458). The efficiency of editing is comparable to the original system (approximately 80%).

**Figure 1.**
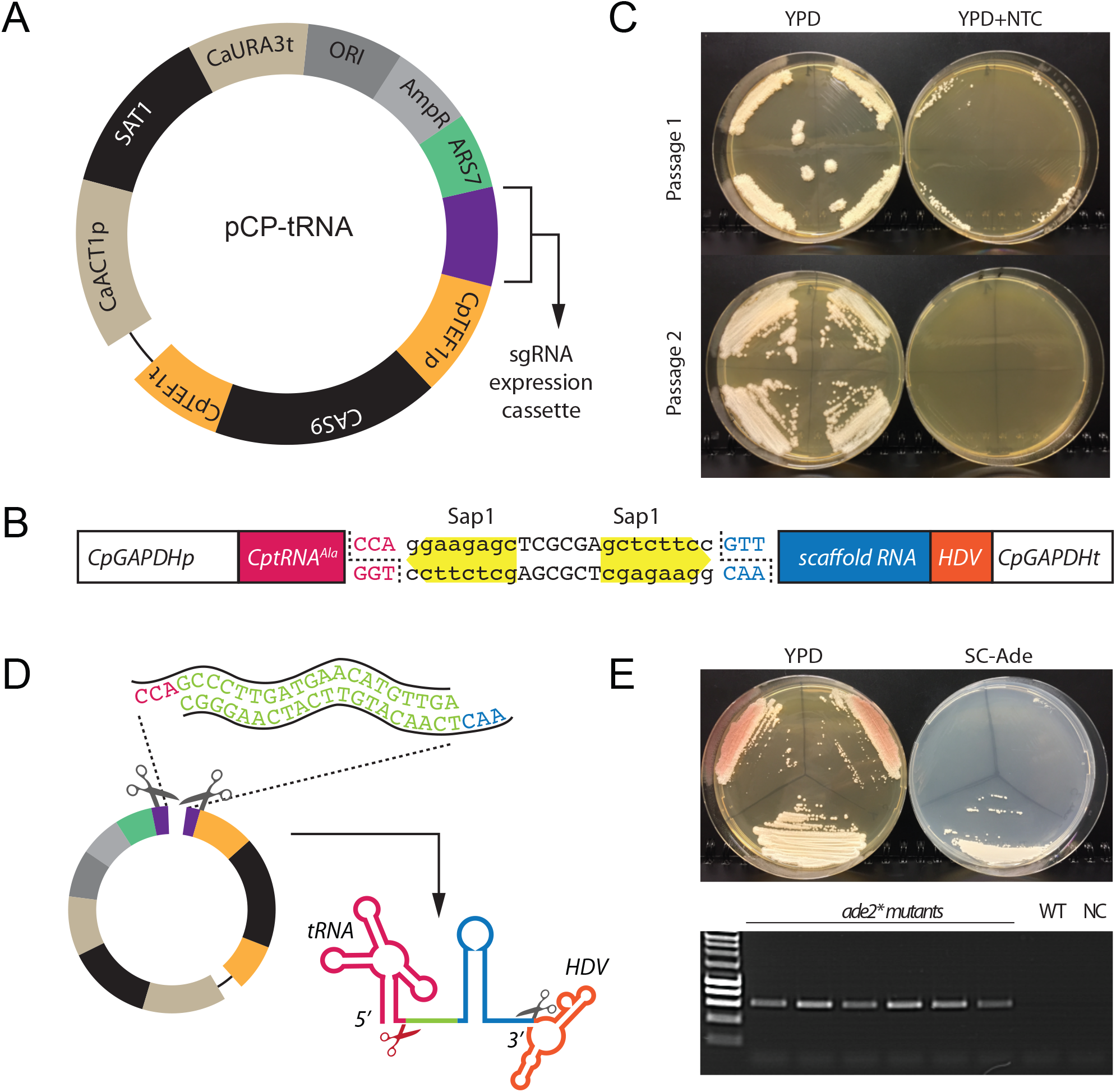
The pCP-tRNA plasmid system for gene editing in *C. parapsilosis*. (**A**) The plasmid shares the main features of the pRIBO system (27), namely the *SAT1* gene (nourseothricin resistance), the autonomously replicating sequence 7 *(ARS7)* from *C. parapsilosis*, and the *CAS9* gene expressed from the *C. parapsilosis TEF1* promoter. (**B**) The pRIBO and pCP-tRNA systems differ in the cassette used to express the sgRNA. In pCP-tRNA, the RNA pol II GAPDH promoter is followed by the tRNA^Ala^ sequence (in pink), two SapI restriction sites (in yellow), the scaffold RNA (in blue), and the Hepatitis Delta Virus (HDV) sequence (in orange). (**C**) Like pRIBO, pCP-tRNA is easily lost. Transformed cells were patched to YPD plates without nourseothricin (NTC) for 48 h, and then streaked on YPD and YPD + NTC. Colonies from YPD were repatched after 48 hr. All transformants lost NTC resistance after just two passages. (**D**) The target guide (in green, ADE2-B in the figure) is generated by annealing two 20 bp oligos carrying overhang ends (in pink and blue), and cloned into SapI-digested pCP-tRNA. The guide RNA is released by cleavage after the tRNA^Ala^ and before the HDV ribozyme. (**E**) Editing of *ADE2* using the pCP-tRNA system. Transformation of *C. parapsilosis* CLIB214 with pCP-tRNA-ADE2-B and a repair templates (RT-B, (27)) resulted in the introduction of two stop codon that disrupted the gene function, producing pink colonies that failed to grow in the absence of adenine (SC-ade). A white Ade+ wild type colony is shown as control. The transformants were screened by PCR using the mutADE2B-F primer derived for the edited site and the downstream ADE2_REV primer, which generates a product only when the mutation is present as described in (27). WT: CLIB214 strain; NC: no DNA.

We next explored the possibility of introducing unique molecular barcodes at the edited site, while retaining short repair templates. We targeted the gene *CPAR2_101060* as proof of principle, using a repair template with 30 bp of homology arms flanking 11 bp containing stop codons in all open reading frames, and a unique tag (Fig. 2A). PCR screening of 15 transformants showed that the barcode was incorporated in all (Fig. 2A). The edited site was confirmed by sequencing. This approach will be useful in large scale studies, as it results in the disruption of the gene regardless of the reading frame, and can be used to specifically barcode each mutant strain for use in competition studies. We also showed that short regions (30 bp) can be used to drive homologous recombination in *C. parapsilosis*.

**Figure 2.**
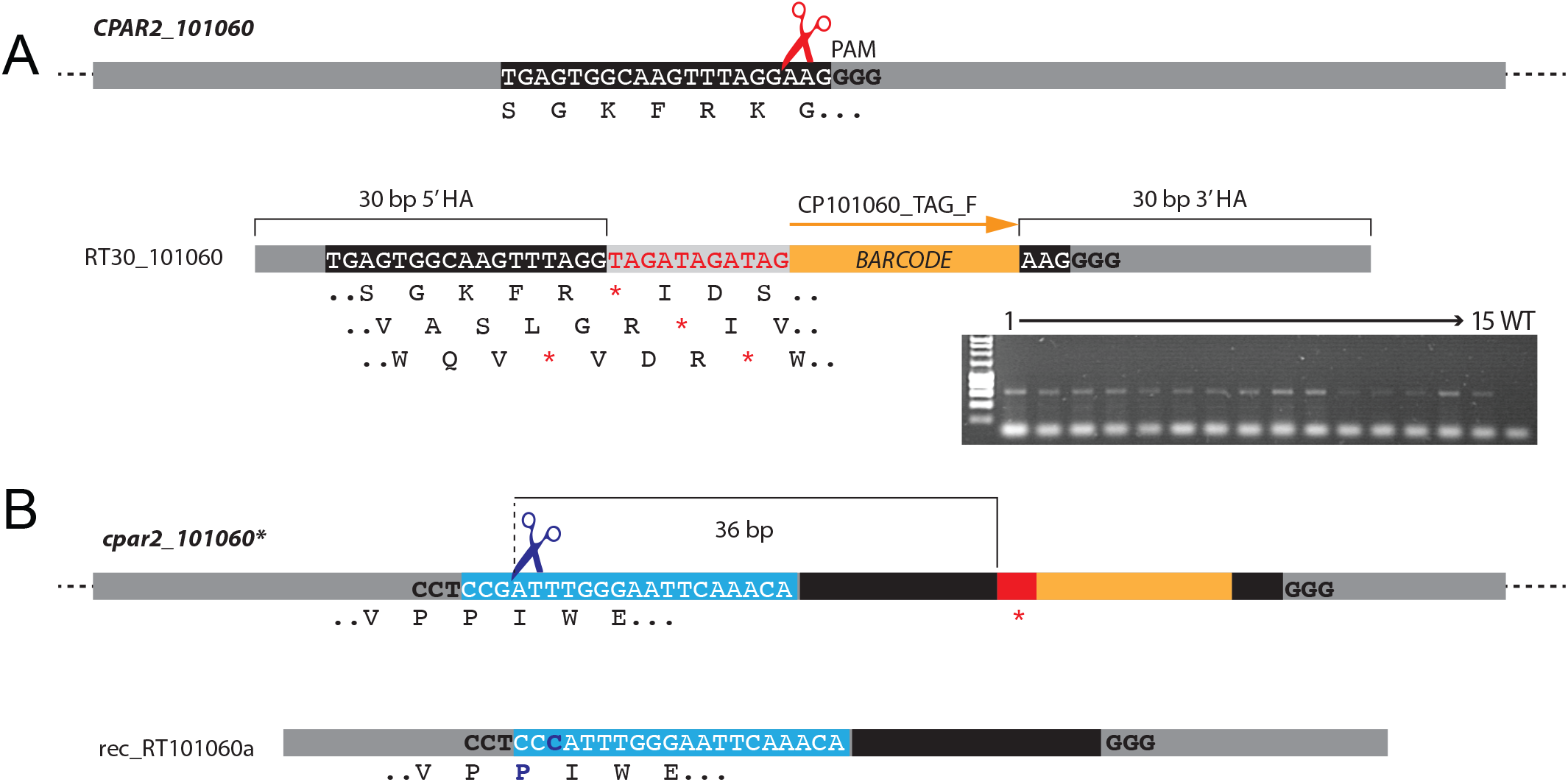
Editing and reconstitution of *CPAR2_101060*. (**A**) The plasmid pCP-tRNA-CP101060 was generated to target *CPAR2_101060*. The guide sequence recognised by Cas9 is boxed in black, and the PAM is shown in bold. The Cas9 cut site is indicated by red scissors. *C. parapsilosis* CLIB214 cells were transformed with pCP-tRNA-CP101060 and a repair template (RT30_101060) generated by overlapping PCR of RT30_101060_TOP and RT30_101060_BOT oligos. The repair template contains two 30 bp homology arms (HA) that flank an 11 bp sequence containing coding stop codons in all three possible reading frame (in red, with all reading frames indicated below), and a 20 bp unique barcode (in orange). The gel shows screening of 15 transformants by PCR using the primer CP101060_TAG_F, which anneals to the barcode, together with the CP101060_WT_R downstream primer. Sequencing confirmed that stop codons were introduced into both alleles of *CPAR2_101060*. (**B**) To replace the *cpar2_101060*\* edited alleles with wild type sequences, a PAM site (bold) upstream from the edited site (red) was selected. The guide RNA is boxed in light blue. Transformation with pCP-rec-tRNA-Cp101060a containing this guide results in Cas9 cleavage 36 bp upstream from the mutated region (indicated by blue scissors). The repair template (rec-RT101060a) generated by overlapping PCR with primers rec-RT-101060aTOP and rec-RT-101060aBOT is designed to replace the edited site and bar code with wild type sequences. It also includes a single G1420C synonymous SNP, so that the reconstituted and wild type alleles can be distinguished. The wild type sequence was successfully reintroduced in 2/9 transformants tested. The scheme is not drawn to scale.

Nguyen et al. (30) showed that CRISPR-Cas9 can be used to reconstitute mutated strains in *C. albicans* by reintroducing the wild type sequence at the native locus. In that study (30), the edited sequence included a new PAM site that was targeted with a common gRNA. To avoid adding even more sequences at the edited site, we instead selected a naturally occuring PAM site 36 bp upstream from the edited region (Fig. 2B). A synonymous SNP (G1420C) was introduced into the repair template so that we could distinguish the reconstituted strain from the original wild type strain (Fig. 2B). From 9 sequenced transformants, two contained the reconstituted wild type sequence, removing the stop codon and the barcode from both alleles, and introducing the synonymous SNP (reconstituted allele: *REC101060*, Table S1, doi:10.6084/m9.figshare.7752458). The remaining 7 all contained the synonymous SNP, but they also retained the edited region and barcode, possibly due to the lower efficiency of homologous recombination with increasing distance from the Cas9 cut site, as discussed below (38).

### CRISPR-Cas9 gene editing in *C. metapsilosis*

Zoppo et al (36) showed that the original plasmid designed for CRISPR editing in *C. parapsilosis* can be used to edit genes in *C. orthopsilosis*, and the tRNA-based plasmid is also effective in this species (Morio et al, in preparation). We therefore tested the pCP-tRNA system in *C. metapsilosis*, the third member of the *C. parapsilosis* species group, by targeting *CmADE2*. Unlike *C. parapsilosis* where heterozygosity levels are low, all *C. metapsilosis* isolates characterized to date descend from hybridization between two parental strains that differ by about 5% at the sequence level (39). The pCP-tRNA plasmid is easily introduced into *C. metapsilosis* SZMC8093, where it propagates without integration. Similar to *C. parapsilosis*, nourseothricin resistance is quickly lost in the absence of selection (Fig. 3A). Cells transformed with a plasmid targeting *CmADE2* and a repair template carrying 35 and 48 bp of upstream and downstream homology regions designed to introduce two stop codons were edited with 100% efficiency. All transformants were pink when replica plated on YPD, and failed to grow on synthetic media in the absence of adenine (Fig. 3B, right plate). PCR analysis of 95 transformants (Fig. S1) using a allele specific primer showed that 88 had the expected mutation, which was confirmed by sequencing 6 representative transformants. Repair by homologous recombination therefore occurs at a high rate in *C. metapsilosis* when a suitable repair template is provided. Interestingly, in the absence of the repair template, most colonies were also pink and adenine auxotrophs. Sequencing of 12 transformants showed that a variety of Non-Homologous End Joining-like repair events had occurred, including deletion of either 2 or 3 nucleotides, and insertion of 1 nucleotide, resulting in either a frameshift or the deletion of one amino acid (Fig. 3D). This suggests that repair via NHEJ is common in *C. metapsilosis* (Fig. 3D).

**Figure 3.**
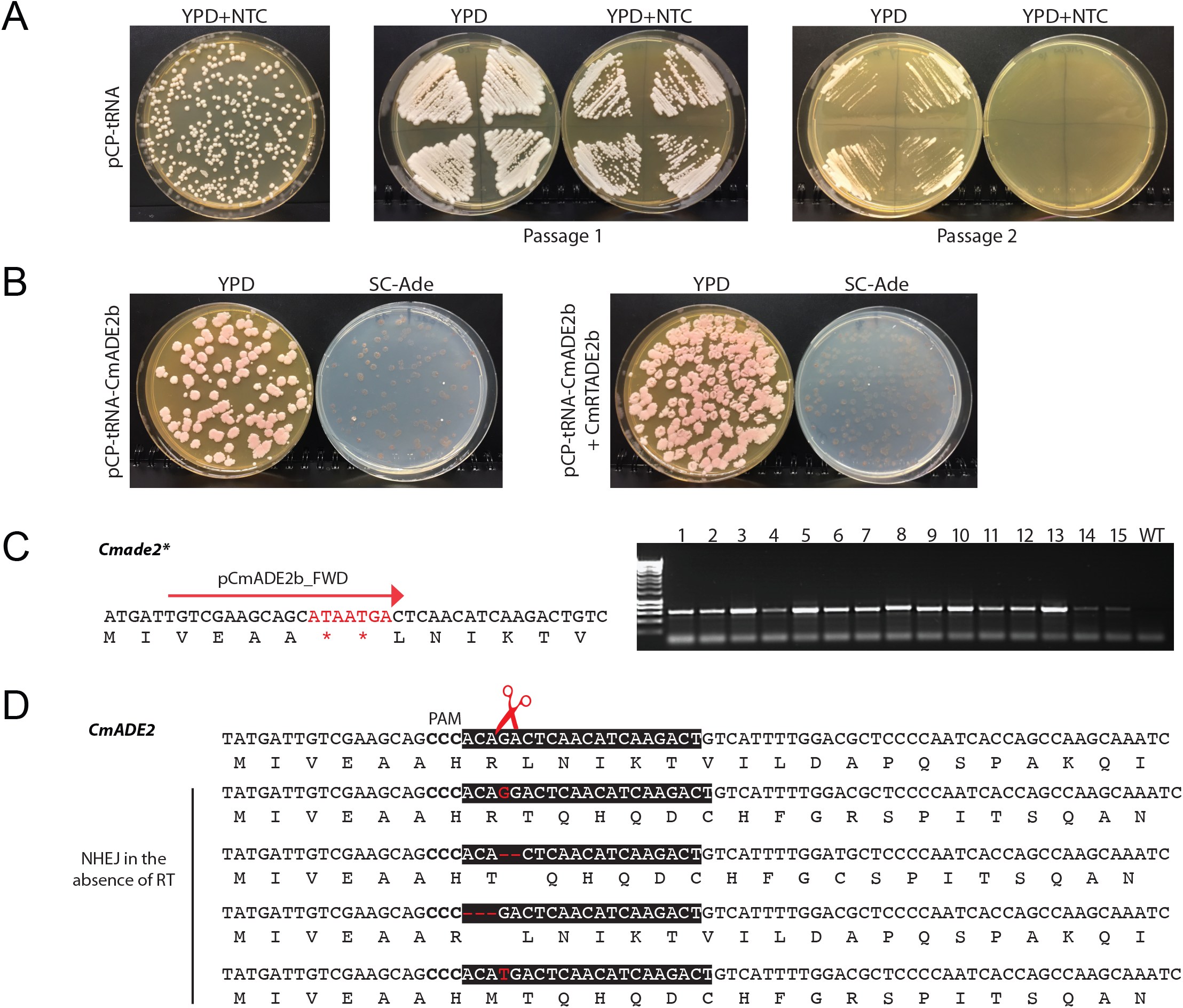
Editing of *ADE2* in *C. metapsilosis*. (**A**) Plasmid pCP-tRNA propagates in *C. metapsilosis* SZMC8093, and it is lost after two passages on YPD in the absence of nourseothricin (NTC, 200 μg/ml). (**B**) *C. metapsilosis* was transformed with plasmid pCP-tRNA-CmADE2b, targeting *CmADE2* either without (left hand side) or with (right hand side) a repair template (CmRTADE2b, generated by overlapping PCR with primers CmRTADE2b_TOP/CmRTADE2b_BOT) designed to introduce two stop codons. The transformants were replica plated on YPD and Sc-Ade. Almost all colonies are pink, and cannot grow in the absence of adenine. (**C**) In the presence of the repair template, most transformants contained the inserted stop codons, identified by PCR using primers pCmADE2b_FWD and CmADE2_REV. Colony PCR of 15 representative colonies is shown, more are shown in Fig. S1 (doi:10.6084/m9.figshare.7752458). The wild type strain (WT) was included as control. (**D**) Many pink transformants were obtained even in the absence of the repair template. Sequencing of the region surrounding the Cas9 cut site revealed a variety of repair events, including insertions and deletions (indicated in red), resulting in either frameshift or deletion of His23. These presumably result from NHEJ.

### Designing a plasmid for gene editing in *C. tropicalis*

Some advantages of using a replicating plasmid-based gene editing system are that any gene can be edited in any isolate in a markerless way. Currently, there are few methods available for editing in *Candida* species other than *C. albicans* and the *C. parapsilosis* species group. We found that the pCP-tRNA plasmid failed to generate nourseothricin resistant transformants of *C. tropicalis*. We therefore adapted constructs designed by Defosse et al (40) who identified suitable promoters and terminators for use in *C. tropicalis*. We first replaced the *GFP* gene in pAYCU268 from Defosse et al (40) with *CAS9* from Vyas et al (25), placing *CAS9* under the control of the *TEF1* promoter from *Meyerozyma guilliermondii*. pAYCU268 expresses *SAT1* from the *C. dubliniensis TEF1* promoter, and is designed to integrate randomly in the *C. tropicalis* genome. We identified an autonomously replicating sequence (CaARS2) which is reported to promote replication in *C. tropicalis* (41, 42), and introduced it into the pAYCU268 backbone. Finally, a tRNA cassette similar to that in pCP-tRNA was synthesized and inserted into the plasmid, generating pCT-tRNA (Fig. 4A). Guide RNAs can then be cloned between the tRNA and a ribozyme, and are expressed from an *Ashbya gossypii TEF1* promoter, followed by the *Saccharomyces cerevisiae CYC1* terminator. An important feature of this plasmid is that, apart from CaARS2, all the DNA parts can be changed thanks to the presence of specific restriction sites.

**Figure 4.**
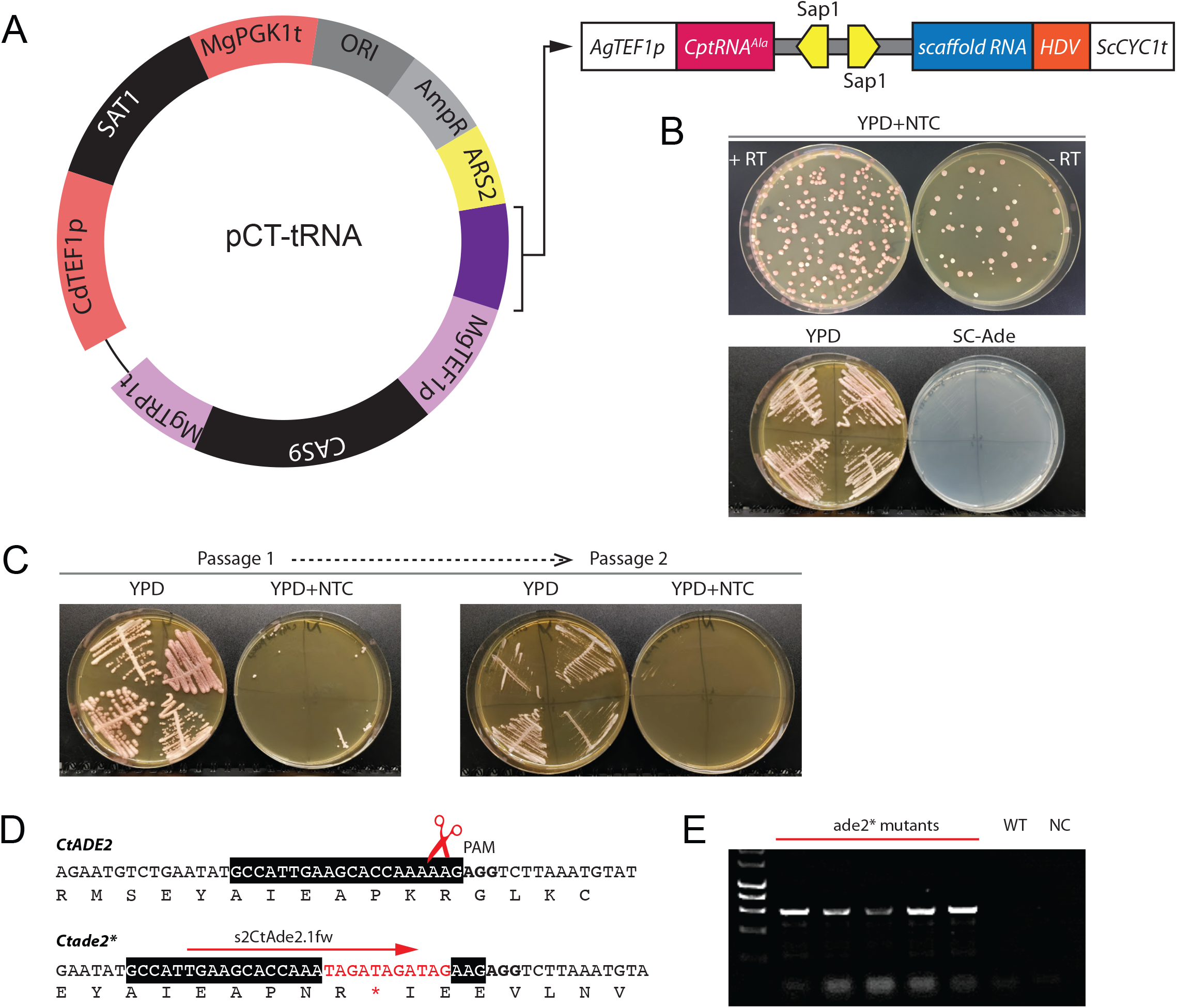
The pCT-tRNA plasmid system for gene editing in *C. tropicalis*. (**A**) The plasmid shares the main features of the pCP-tRNA system, except that different regulatory elements and a different autonomously replicating sequence is used. The *SAT1* gene (nourseothricin resistance) is flanked by a *CdTEF1* promoter and a *MgPGK1* terminator, and the *CAS9* gene is expressed from the *M. guilliermondii TEF1* promoter. The autonomously replicating sequence 2 (*ARS2*) was derived from *C. albicans* (41, 42). The cassette for the expression of the sgRNA is highlighted in purple on the plasmid map, and represented in more details in the scheme on the right side. The color coding is the same as in Fig. 1. The only differences with the cassette in Fig. 1B are the promoter *(AgTEF1p)* and the terminator (*ScCYC1t*). (**B**) Editing of *CtADE2* using the pCT-tRNA system. The guide gCtADE2.1 was generated by annealing CtAde2.1_gTOP and CtAde2.1_gBOT oligonucleotides (Table S2, doi:10.6084/m9.figshare.7752458), and cloned into the SapI-digested plasmid pCT-tRNA to generate pCT-tRNA-CtADE2.1. Transformation of *C. tropicalis* Ct46 with pCT-tRNA-CtADE2.1 and the repair template R60-CtADE2-b resulted in the introduction of a stop codon that disrupted the gene function, producing pink auxotrophs. Pink colonies were also observed when cells were transformed with pCT-tRNA-CtADE2.1 without any repair template, presumably due to NHEJ-like repair events (see also Fig. S3 (doi:10.6084/m9.figshare.7752458)). (**C**) pCT-tRNA-CtADE2.1 is easily lost. Representative pink colonies were patched to YPD plates without nourseothricin (NTC) for 48 h, and then streaked on YPD and YPD + NTC. Colonies from YPD were repatched after 48 hr. All transformants lost NTC resistance after just two passages. (**D**) The transformants were screened by PCR using the s2CtAde2.1fw primer derived from the edited site and the downstream pCtADE2.1_REV primer, which generates a product only when the mutation is present. (**E**) Result of PCR screening of 5 representative transformants. WT: Ct46 strain; NC: no DNA.

The pCT-tRNA plasmid containing a guide RNA targeted against *CtADE2* was used to transform 5 different isolates of *C. tropicalis*, together with a repair template designed to introduce one stop codon in each frame (Fig. 4D, Fig. S3 (doi:10.6084/m9.figshare.7752458)). The repair template has 60 bp homology arms flanking the cleavage site, and was generated by overlapping PCR. Pink auxotrophs were observed following transformation of each isolate, in the presence or absence of repair template, with an efficiency ranging from 88% to 100% (Fig. 4B, Table 1). PCR screening and subsequent sequencing analysis of representative transformants obtained using the repair template confirmed the presence of the inserted stop codon (Fig. 4D and E). In the absence of the repair template, pink adenine auxotrophs resulted from the deletion of one base near the Cas9 cut site (Fig. 4B, Fig. S3A, Table 1). Non-Homologous End Joining-like repair events therefore occur at a high frequency in *C. tropicalis*. Just like pCP-tRNA, the pCT-tRNA plasmid is rapidly lost from transformants grown in the absence of selection (Fig. 4), this minimizing the risk of Cas9 cutting at off-target sites.

**Table 1.**
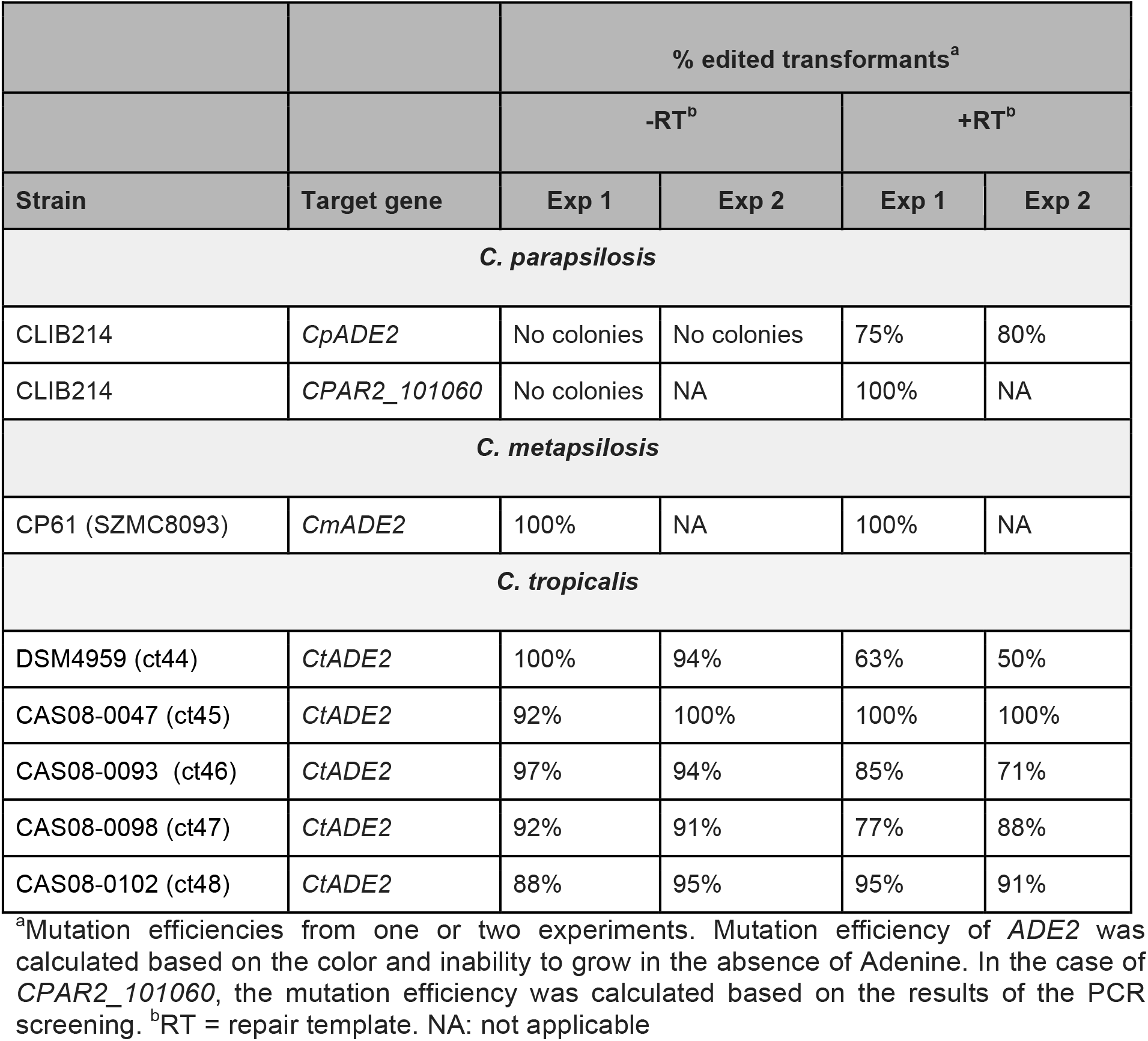
Efficiency of CRISPR-based editing with the pCP/CT-tRNA systems in *C. parapsilosis, C. metapsilosis* and *C. tropicalis*.

### Using CRISPR to introduce heterozygous mutations

One of the problems with CRISPR editing of diploid genomes is that it tends to be an all-or-nothing affair - either both alleles are edited, or neither allele is. This can be a problem with essential genes, where editing both alleles would be lethal. Vyas et al (24) approached this problem by generating temperature sensitive alleles in *C. albicans*. However, Paquet and colleagues (38) recently described how the design of the repair template can be used to introduce mutations into just one allele in the genome of human cells. They showed that there is an inverse relationship between the efficiency of incorporation of a desired SNP with its distance from the Cas9-induced double strand break. This can be exploited to push the editing system towards the introduction of mutations at one allele only. Alternatively, cells transformed with a mixture of two different repair templates can incorporate a different template at each allele (38). We tested if the same approaches are effective in *C. parapsilosis* by targeting the (non-essential) gene *CPAR2_101060*.

Both strategies were used. In strategy 1, *C. parapsilosis* CLIB214 was transformed with a plasmid targeting *CPAR2_101060*, and one of two repair templates. Both repair templates carry three base substitutions (GGG>ATC) that disrupt the PAM, and introduce an Ile154Gly amino acid change. In addition, one repair template contains one synonymous SNP (C>G) 10 bp upstream from the Cas9 cut site, and the second contains a different synonymous SNP (T>C) 19 bp upstream from the cut site (Fig. 5A, RTs Het_LL1 and Het_LL2, respectively). All the transformants obtained with either repair template that were tested by PCR contained the ATC amino acid change that also disrupts the PAM (Fig. 5A). Seven transformants obtained with the first repair template (Het_LL1) were all homozygous (either C or G) at the position 10 bp upstream from the cut site (Fig. 5A, upper panel). Using the second repair template (Het_LL2), 2 of 6 transformants tested were homozygous for the wild type nucleotide, 3 incorporated C at both alleles, and one was heterozygous, with T at one allele and C at the second (Fig. 5A, lower panel). Increasing the mutation-to-cut site distance can therefore increase the chances of obtaining heterozygous substitutions in *C. parapsilosis*. It is probably necessary to disrupt the PAM site or the protospacer when using this strategy, to prevent repeated recutting by Cas9.

**Figure 5.**
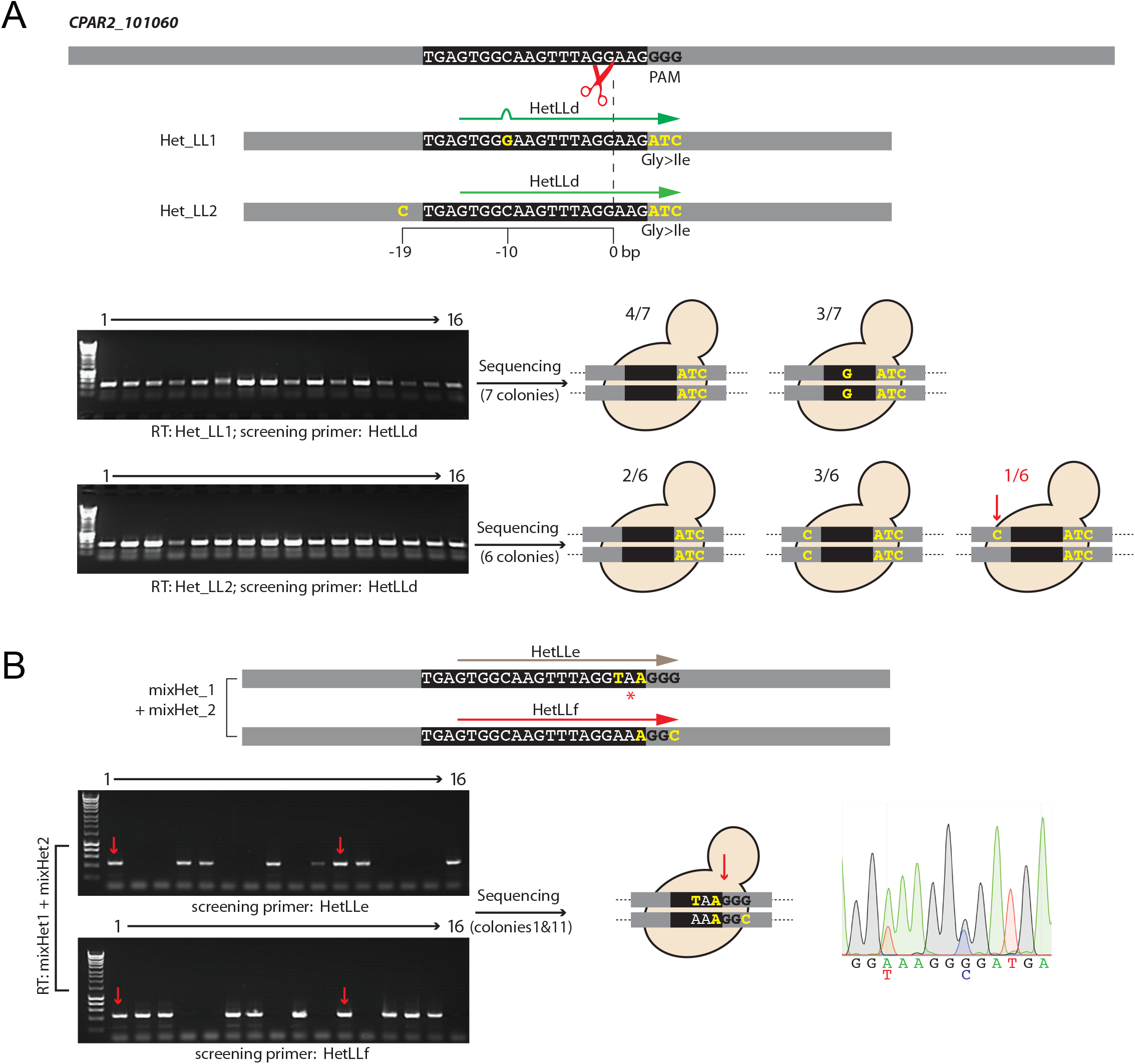
Generation of heterozygous mutations with the CRISPR-Cas9 system. (A) Varying the distance between the Cas9 cut site and introduced variation, (B) Using a mixture of repair templates. (**A**) *C. parapsilosis* CLIB214 cells were transformed with pCP-tRNA-CP101060 and either the repair template Het_LL1 or the repair template Het_LL2. Both Het_LL1 and Het_LL2 carry three nucleotide changes that introduce a codon change and disrupt the PAM (GGG to ATC, resulting in a Gly to Ile amino acid change) (yellow). The two templates also carry an additional SNP which introduces a silent mutation, either 10 bp (C>G) or 19 bp (T>C) upstream from the Cas9 cut site. Sixteen transformants obtained with Het_LL1 and 16 transformed with Het_LL2 yielded a PCR product using primers HetLLd which anneals at the ATC codon change, and the CP101060_WT_R downstream primer. HetLLd does not anneal perfectly to the sequence introduced by Het_LL1, because of the C>G SNP. The cartoons show the sequencing results. With repair template Het_LL1, all 7 transformants contained the amino acid change at both alleles, in three G was introduced at both alleles 10 bp upstream of the cut site, and four retained the wildtype C base. With repair template Het_LL2, again all 6 transformants contained the amino acid change at both alleles, 5 contained either the wildtype or the mutated base at the additional site, but one was heterozygous for T>C SNP 19 bp from the cut site. The combination of alleles in this strain is referred to as *CP101060-ATC/CP101060-ATC-SNP*. (**B**) *C. parapsilosis* CLIB214 was transformed with with pCP-tRNA-CP101060 and a mixture of two repair templates, mixHet_1 and mixHet_2. MixHet_2 is designed to introduce two silent mutations that do not change the coding sequence, but that do disrupt the PAM site, preventing Cas9 from cutting again at the the edited site. mixHet_1 also contains two mutations, which changes the protospacer preventing the gRNA binding, and which introduce a stop codon. From sixteen transformants, two (indicated by the red arrows), generated PCR products when amplified with primers annealing to either of the targeted editing sites (HetLLe or HetLLf) and the CP101060_WT_R downstream primer. Sequencing of these colonies confirmed that different repair templates had been incorporated at each allele. The chromatogram shows one example. Genotype: *CP101060-Stop/CP101060-SNP*.

In strategy 2, two repair templates (mixHet_1 and mixHet_2, Fig. 5B) were supplied simultaneously. Both were designed to introduce synonymous SNPs that disrupt either the PAM or the protospacer. One repair template also contains a stop codon immediately upstream of the PAM. Incorporation of one repair template at one allele and the other at the second allele should generate a functional heterozygote, in which one allele has a stop codon and one does not. PCR screening showed that one or other repair template was incorporated in 14 of 16 transformants tested (Fig. 5B). For two colonies, PCR and sequence analysis showed that both repair templates were incorporated, with a stop codon introduced at only one allele (Fig. 5B). The presence of the synonymous SNPs at both alleles indicated that homology-directed repair had occurred at both, one using repair template 1, and one repair template 2. Two different strategies can therefore be used in combination with the pCP-tRNA plasmid to introduce heterozygous mutations.

## Conclusion

The pCP/CT-tRNA systems can be used for CRISPR-Cas9 mediated gene editing in all three members of the *C. parapsilosis sensu latu* complex, and in *C. tropicalis*. CRISPR editing can also be used to reconstitute wild type alleles, and to generate heterozygous mutations.

## Methods

### Strains and media

All *C. parapsilosis, C. metapsilosis*, and *C. tropicalis* strains used in this study (Supplementary Table S1, doi:10.6084/m9.figshare.7752458) were grown in YPD medium (1% yeast extract, 2% peptone, 2% glucose) or on YPD plates (YPD + 2% agar) at 30 °C. Transformants were selected on YPD agar supplemented with 200 μg/ml nourseothricin (Werner Bioagents Jena, Germany). Auxotrophies were confirmed by growing mutant strains on synthetic complete dropout media (0.19% yeast nitrogen base without amino acids and ammonium sulfate, 0.5% ammonium sulfate, 2% glucose, 0.075% amino acid dropout mix, 2% agar). All the plasmids used in this study (Supplementary Table S3, doi:10.6084/m9.figshare.7752458) were propagated in *Escherichia coli* DH5α cells (NEB, UK) by growing cells in LB media without NaCl (Formedium) supplemented with 100 μg/ml Ampicillin (Sigma).

### Construction of the pCP-tRNA series plasmids

The synthetic construct GAPDHp-tRNA-SapI-HDV (Eurofins MWG, Fig. S2 (doi:10.6084/m9.figshare.7752458)) was designed to include the *C. parapsilosis GAPDH (CPAR2_808670)* promoter, a *C. parapsilosis* tRNA^Ala^ sequence, two tandem SapI/BspQI sites for the cloning of the guide, a Hepatitis Delta virus (HDV) ribozyme, and the *GAPDH* terminator. The cassette was cloned by Gibson Assembly (primers in Table S2, doi:10.6084/m9.figshare.7752458) into NruI-digested pSAT3 plasmid, which differs from the published pSAT1 plasmid (27) only in that it does not contain any SapI/BspQI sites. The guide RNA was generated by annealing of two 23 bp oligos carrying appropriate overhanging ends, and cloned into the SapI-digested pCP-tRNA plasmid (Supplementary Tables S2 and S3).

### Construction of pCT-tRNA plasmids

pAYCU268, from Defosse et al (40) was used as a starting vector. The *GFP* gene was replaced with *CAS9* from pV1326 (25) by Gibson assembly using primers jpGA_pAYCU268.fw and GA_pAYCU.CAS9.rv to amplify the backbone and the jpGA_CAS9.1326.fw and pGA_Cas9.1326.rv to amplify *CAS9*. resulting the plasmid named GA_pAYCU.CAS9.1326. CaARS2 (41) was amplified using V3_GA.CaARS2.1326.fw and V3_GA.CaARS2.1326.rv, and introduced into the backbone amplified with the primers GA_pAYCUCAS9.1326.fw and GA_pAYCUCAS9.1326.rv by Gibson assembly. A SapI site in the plasmid backbone was removed in this step. A synthetic construct (Eurofins MWG, Fig. S2 (doi:10.6084/m9.figshare.7752458)) was designed as described for pCP-tRNA, except that the expression of the sgRNA is driven from the *A. gossypii TEF1* promoter and *CYC1* terminator from S. *cerevisiae*. The AgeI and SpeI restriction sites flanking the cassette were used for cloning into the double digested GA_pAYCUCAS9.1326.ARS vector. *C. tropicalis* guide RNAs were designed using CHOPCHOP v2 (43, 44).

### Transformation of *Candida* strains

*C. parapsilosis* CLIB214 and *C. metapsilosis* CP61 strains were transformed using the lithium acetate method as described in (27), with minor modifications. Each repair template was generated by primer extension of overlapping oligonucleotides (Supplementary Table S1, Table S2, doi:10.6084/m9.figshare.7752458) and 25 μl of unpurified product were used to transform yeast cells. Transformation of *C. tropicalis* strains was performed by using a modified electroporation protocol (12, 45, 46). *C. tropicalis* cells were grown to A_600_ of 5-10 and then resuspended in 0.1 M lithium acetate, 10 mM Tris·HCl pH 8.0, 1 mM EDTA, 10 mM DTT and incubated at room temperature for 1 h. Cells were washed twice with ice cold water and once in 1 M ice cold sorbitol. The sorbitol wash was decanted, and cells were resuspended in the remaining liquid. Approximately 40-50 μl of cells were used per transformation, with 5 μg of plasmid together with 5 μg of purified repair template. Cells and DNA were electroporated at 1.8 kV by using a Bio-Rad Pulser Xcell™ Electroporator and immediately resuspended in 1 ml of cold 1 M sorbitol. Cells were subsequently resuspended in 1 ml of YPD and allowed to recover for 4 h at 30 °C before plating on selective media (YPD + 200 μg/mL NAT). Nourseothricin-resistant transformants were patched onto YPD (and SC lacking adenine where indicated), and screened by colony PCR. The mutation efficiency was calculated as (edited transformants on the plate)*100/(total number of transformants on the plate). Representative mutants were sequenced by Sanger sequencing (Eurofins MWG). Loss of the plasmid was induced by patching transformants onto YPD agar without selection and re-patching every 48 h until they no longer grew on YPD agar plates containing 200 μg/ml nourseothricin.

## Acknowledgements

We are grateful to Prof N. Papon from Université Bretagne-Loire for the gift of plasmids. This work was supported by awards from Science Foundation Ireland (12/IA/1343, https://www.sfi.ie) and the European Union’s Horizon 2020 research and innovation program under the Marie Sklodowska-Curie grant agreement no. H2020-MSCA-ITN-2014-642095.

## Supplementary materials are available at FigShare, doi:10.6084/m9.figshare.7752458

**Figure S1**. PCR screening of *C. metapsilosis* SZMC8093 transformants.

Cells transformed with pCP-tRNA-CmADE2b + CmRTADE2b were screened with pCmADE2b_FWD and CmADE2_REV.

**Figure S2**. Sequence of the synthetic tRNA cassettes.

(A) GAPDHp-tRNA-SapI-HDV construct for pCP-tRNA.

(B) AgTEF1p-tRNA-SapI-HDV-ScCYC1t construct for pCT-tRNA.

**Figure S3**. *CtADE2* editing in *C. tropicalis* isolates. (A) Transformants of *C. tropicalis* growing on YPD + NTC plates. The strains are indicated above each picture. +RT = cells transformed with pCT-tRNA-ADE2.1 + R60-CtADE2-b. -RT = cells transformed with pCT-tRNA-ADE2.1. (B) The wildtype sequence of *CtADE2* is shown. The guide sequence is boxed in black. Red scissors indicate the cut site. Representative colonies from strains Ct44, Ct45, Ct46, and Ct47 transformed without the RT were sequenced. Non-Homologous End Joining-like repair events resulted in the deletion of one A in proximity of the cut site in all the colonies tested.

**Table S1**. Strains used/generated in this study

**Table S2**. Oligonucleotide sequences used in this study

**Table S3**. Plasmids generated in this study

